# Whole-night gentle rocking improves sleep in poor sleepers with insomnia complaints

**DOI:** 10.1101/2025.03.31.646264

**Authors:** Aurore A. Perrault, Nathan E. Cross, Thien Thanh Dang Vu, Sophie Schwartz, Laurence Bayer

**Affiliations:** Department of Neuroscience, Faculty of Medicine, University of Geneva, Geneva, Switzerland; Department of Health, Kinesiology and Applied Physiology, Concordia University, Montreal, Canada; Centre de Recherche de l’Institut Universitaire de Gériatrie de Montréal, CIUSSS Centre-Sud-de-l’Ile-de-Montréal, Montreal, Canada; Sleep & Circadian Research Group, Woolcock Institute of Medical Research, Macquarie University, Sydney, NSW, Australia; Brain and Mind Centre, School of Psychology, The University of Sydney, NSW, Australia; Swiss Center for Affective Science, Campus Biotech, Geneva, Switzerland; Center for Sleep Medicine, Geneva University Hospital, Geneva, Switzerland

**Keywords:** rocking stimulation, insomnia, poor sleep, spindle, slow oscillations, entrainment

## Abstract

Specific brain oscillations can be manipulated during sleep to improve sleep quality and memory performance. We previously demonstrated that continuous rocking stimulation (0.25Hz, lateral movement) applied to good sleepers during sleep enhanced stable deep sleep, boosted NREM oscillations (spindles and slow waves), and memory consolidation. Here, we investigated whether nocturnal rocking could benefit individuals suffering from sleep difficulties. We recruited sixteen young adults with subjective difficulties initiating and/or maintaining sleep and who presented with objective poor sleep quality. Each participant spent two nights of sleep at the laboratory, one rocking and one stationary, during which we assessed sleep and declarative memory consolidation. We found that a whole night of gentle rocking in individuals with poor sleep decreased sleep fragmentation, time spent awake and in light sleep (N1), with an associated increase in objective sleep efficiency and subjective sleep quality. Additionally, we replicated the neural entrainment or synchronizing effect of the rocking motion, yielding a boost in NREM fast spindles and slow oscillations. Yet, these changes in sleep did not modulate overnight memory performance. By alleviating some difficulties encountered in this population of poor sleepers (e.g., sleep maintenance and poor self-reported sleep), these findings provide preliminary evidence that rocking may represent an alternative or complementary intervention for the management of some forms of chronic insomnia.

**STATEMENT OF SIGNIFICANCE:** Here, we demonstrate that a gentle and continuous rhythmic rocking stimulation applied during a whole night improves sleep in young adults with sleep complaints and objective poor sleep quality. Rocking, compared to a ‘normal’ stationary condition, promoted sleep maintenance and sleep efficiency with a parallel improvement in subjective sleep quality. As we previously found in healthy controls, the rocking stimulation had a mechanistic influence over the synchronisation of sleep oscillations in these individuals with insomnia complaints. These findings may be relevant for the development of non-pharmacological interventions in similar populations with insomnia complaints and poor sleep, including older adults, and clinical populations with neurological, psychiatric, or somatic conditions.

## BACKGROUND

We all know that a bad night of sleep is distressing and has negative impacts on our daytime functioning. Normal sleep is composed of an orderly succession of brain states characterized by distinct oscillatory patterns of neural activity, visible on an electroencephalographic (EEG) recording. Specific brain oscillations can be manipulated during sleep to improve sleep quality, memory performance and mental health^1–5^. Recently, we demonstrated that, when healthy young adults spend a nap^6^ or a whole night^7^ on a gently rocking bed (0.25Hz, lateral movement), they slept better, with faster entrance into sleep, prolonged time spent in deep sleep, and fewer nighttime awakenings, while also showing a parallel increase in overnight declarative memory consolidation^7^. We observed similar beneficial effects of rocking on sleep architecture in mice^8^ and demonstrated that these effects were mediated by the otolithic organ of the vestibular system, which sends direct and indirect inputs to the thalamus^8^. The rhythmic stimulation of the vestibular system induced by continuous rocking would thus entrain thalamocortical circuits involved in the generation of electrical sleep brain rhythms, such as sleep spindles and SOs^7,9^.

By specifically targeting sleep entrance and maintenance, sleeping in a rocking bed may benefit individuals suffering from sleep difficulties, such as insomnia disorder. Defined by self-reported complaints of difficulty falling asleep and/or maintaining sleep for more than 3 months, insomnia is accompanied by daytime functioning complaints, such as fatigue, mood disruption and cognitive disturbances^10,11^. Insomnia affects more than 10% of the population^12,13^ and is associated with significant physical and mental health consequences^14–17^. The first line of treatment for chronic insomnia is cognitive-behavioural therapy for insomnia (CBTi)^18,19^, a multimodal psychological intervention aimed at modifying maladaptive thinking and behaviours that contribute to the perpetuation of insomnia^20^. Overall, CBTi reduces insomnia severity by improving self-reported sleep quality (e.g., questionnaires, sleep diaries) and reducing sleep misperception (i.e., mismatch between sleep subjectively reported and objectively recorded)^21^ without changing the objectively recorded sleep architecture^22–24^. Despite high response (>50%) and remission rate (>35%), insomnia symptoms persist in about half of the patients after CBTi^23^. This could be due to the important heterogeneity of the disorder, which encompasses a variety of symptoms, etiologies, and sub-types of insomnia^25–29^. Indeed, as CBTi mainly targets the core subjective symptoms of insomnia, individuals combining psychological and physiological factors (previously called psychophysiological insomnia^30,31^) could benefit from a complementary intervention targeting objective sleep disturbances.

Here, we tested whether these individuals may benefit from the rocking bed which appears to specifically modulate and benefit neurophysiological factors typically associated with insomnia, such as longer sleep latency and time spent in light sleep, as well frequent awakenings. We thus recruited 16 young adults with subjective complaints of difficulties initiating and/or maintaining sleep and presenting with objective poor sleep quality. We recorded their sleep while they spent one night on a rocking bed (0.25 Hz, 10.5 cm lateral excursion)^6,7^ compared to one night in a stationary position (**Figure 1**). We tested whether rocking modulated (i) self-reported sleep quality, (ii) sleep architecture and arousal, and (iii) sleep brain oscillations (i.e., sleep spindles and slow oscillations). Based on our previous work^7^, we also assessed whether rocking and its potential associated effects on sleep may alleviate declarative memory impairments frequently reported in chronic insomnia^32,33^.

**Figure 1.**
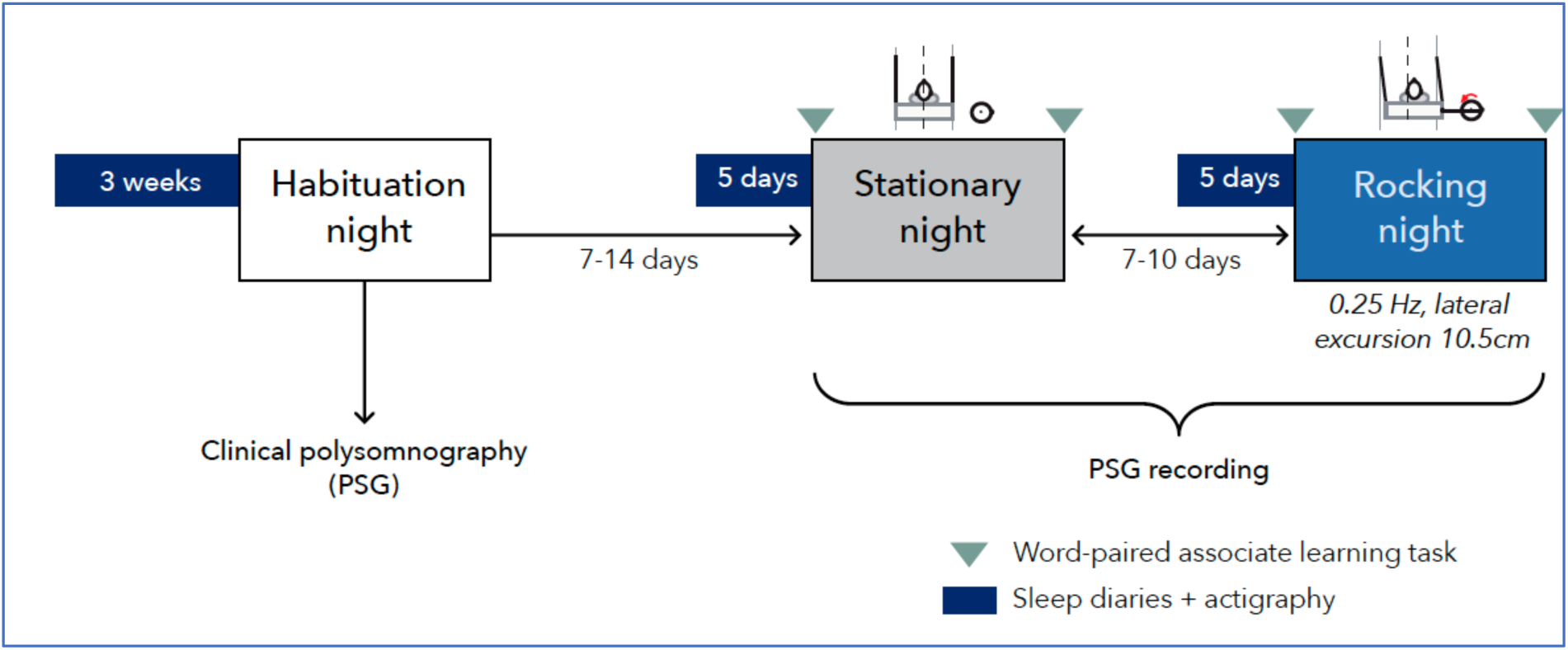
Study design. After 3-week of actigraphy and sleep diary followed by one habituation night, 16 participants underwent two experimental nights: a stationary night (grey), during which the motor (that was used to put the bed into motion) was switched on but not connected to the bed, and a rocking night (blue), during which the bed moved gently (0.25 Hz, 10.5 cm lateral excursion). Both experimental conditions were administered in a randomized order across participants, separated by at least 7 days. Each experimental night was preceded by 5 days of actigraphy and sleep diary. The memory task (word-paired task) was administered in the evening and the morning of both experimental nights.

## METHODS

### Participants

Participants were recruited through print advertisements posted in the Geneva area (Switzerland). Prospective participants were initially screened through online questionnaires for complaints of difficulties falling asleep and/or maintaining sleep and/or early awakening with daytime impairment for at least 3 months. They first filled out the Insomnia Severity Index (ISI)^34^, the Kessler Psychological Distress Scale (K6)^35^, and questions about the use of sleep-inducing substance use. Individuals reporting at least one “severe” sleep difficulty (i.e., sleep entrance, sleep maintenance, early awakening) and an ISI score >8^34^, with no sleep-related substance use and a K6 score <18^36^, were contacted and asked to fill out a series of questionnaires assessing their demographic, medical history, emotional state and sleep habits. Exclusion criteria were as follows: being less than 18 years old and over 35 years old, history or current presence of medical or psychiatric condition (e.g., bipolar disorder, psychosis, cancer), diagnosis of other sleep disorders (e.g., central disorders of hypersomnolence, restless leg syndrome)^10^, moderate to severe anxiety/depression at the Hospital Anxiety and Depression Scale (score >11)^37,38^, and having performed shift work or changed time zones over the past 2 months. Potentially eligible participants subsequently wore a wrist actigraph (Actimeter GT3X+, Actigraph, Pensacola, FL, USA) for 3 weeks and underwent a screening polysomnographic (PSG) recording to rule out the presence of sleep apnea (apnea-hypopnea index >5/h), periodic limb movement (>15/h) and/or cardiac arrhythmia. We also assessed the presence of objective poor sleep (i.e., mean sleep efficiency ≤85%, extracted by 3-week actigraphy and by PSG during the habituation night)^19,39^.

All participants signed a written informed consent form before entering the study, which was approved by the ethics review board of the Geneva University Hospital (Swissethics Committees on research involving humans, CCER Geneva, Switzerland). A total of 21 participants out of 153 participants screened were found eligible. Out of the 21 initially included, one participant voluntarily dropped out after the clinical PSG and 4 were excluded for technical reasons (electrical failure or poor EEG signal). Finally, the sleep of 16 young adults between 18 and 32 years old (11 female) was analysed (see **Table 1** for demographic information).

**Table 1.**
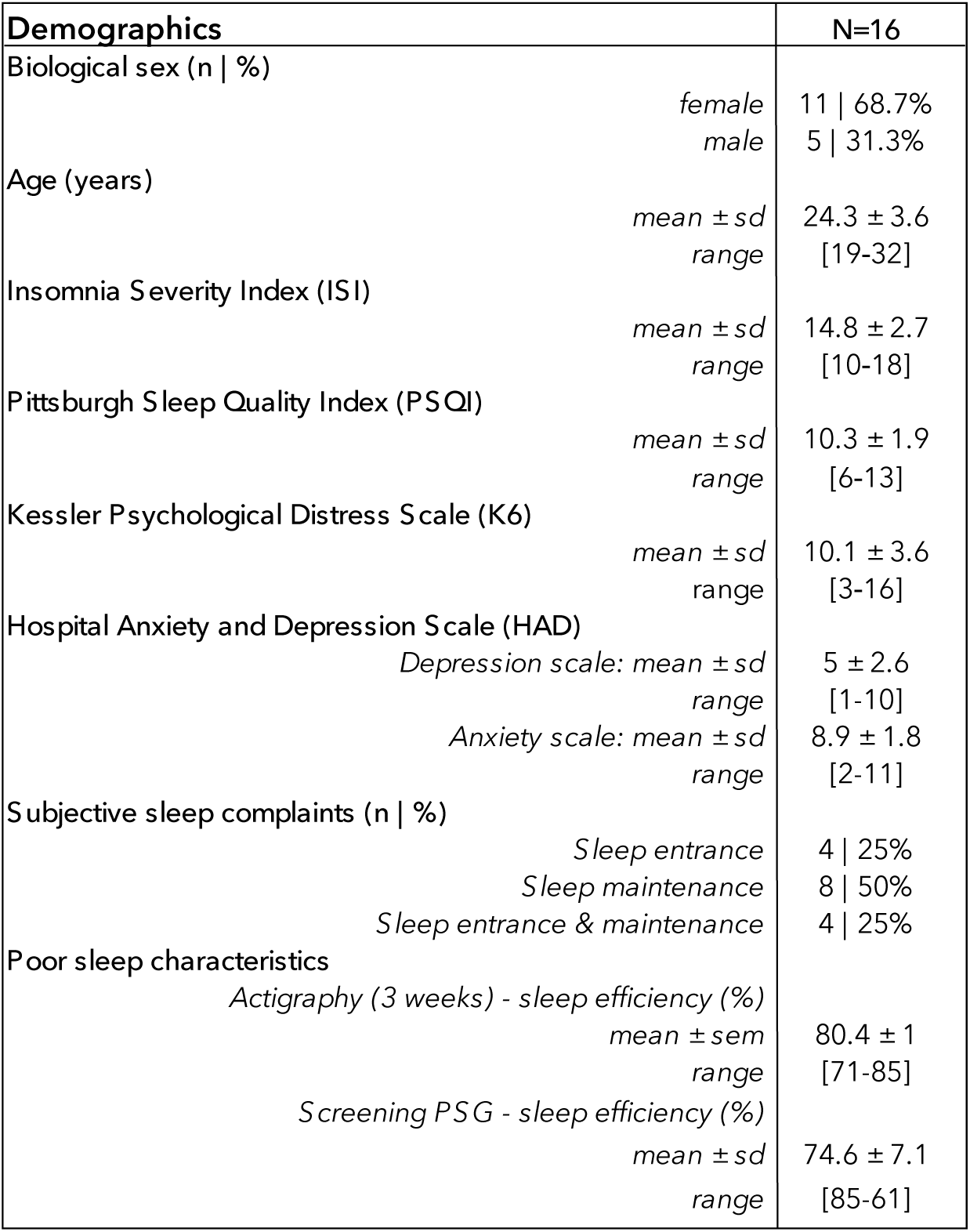
Demographics (N=16)

| Demographics |  | N=16 |
| --- | --- | --- |
| Biological sex (n \| %) | *female* | 11 \| 68.7% |
|  | *male* | 5 \| 31.3% |
| Age (years) | *mean ± sd* | 24.3 ± 3.6 |
|  | *range* | [19-32] |
| Insomnia Severity Index (ISI) | *mean ± sd* | 14.8 ± 2.7 |
|  | *range* | [10-18] |
| Pittsburgh Sleep Quality Index (PSQI) | *mean ± sd* | 10.3 ± 1.9 |
|  | *range* | [6-13] |
| Kessler Psychological Distress Scale (K6) | *mean ± sd* | 10.1 ± 3.6 |
|  | *range* | [3-16] |
| Hospital Anxiety and Depression Scale (HAD) | *Depression scale: mean ± sd* | 5 ± 2.6 |
|  | *range* | [1-10] |
|  | *Anxiety scale: mean ± sd* | 8.9 ± 1.8 |
|  | *range* | [2-11] |
| Subjective sleep complaints (n \| %) | *Sleep entrance* | 4 \| 25% |
|  | *Sleep maintenance* | 8 \| 50% |
|  | *Sleep entrance & maintenance* | 4 \| 25% |
| Poor sleep characteristics | *Actigraphy (3 weeks) - sleep efficiency (%)* |  |
|  | *mean ± sem* | 80.4 ± 1 |
|  | *range* | [71-85] |
|  | *Screening PSG - sleep efficiency (%)* |  |
|  | *mean ± sd* | 74.6 ± 7.1 |
|  | *range* | [85-61] |

### Protocol

Eligible participants followed the same experimental procedure and slept in the same rocking apparatus as described previously^6,7^ (see **Figure 1A**). To summarize, within a month after the initial clinical and habituation PSG, participants came back to the sleep laboratory and spent two experimental nights under PSG monitoring: one with the bed in a stationary position and one with the bed rocking gently (rocking lateral excursion of 10.5cm, frequency of 0.25Hz, i.e., 4s to complete a full back and forth excursion) during the whole night. The order of the two conditions (stationary, rocking) was counterbalanced across participants and separated by a minimum of 7 days. For both experimental sessions, participants performed a declarative memory task (word paired-associate learning task)^7^ in the evening and the morning. After each night, participants filled out questionnaires about the subjective quality of their sleep and rocking pleasantness (specifically after the rocking night).

## Measures

### Polysomnographic (PSG) recording

Whole-night PSG recording was used for the habituation screening and the experimental nights. PSG included EEG, EOG, EMG, ECG and a thoracic belt, recorded with a Deltamed amplifier (Natus Europe, GmbH, Germany). Twenty-two scalp electrodes (Fp1, Fpz, Fp2, F7, F3, Fz, F4, F8, T3, C3, Cz, C4, T4, T5, P3, Pz, P4, T6, O1, O2, A1, A2) were placed according to the international 10-20 system and referenced to PFz. Breathing measurements were added during the habituation night. The EEG signal was filtered between 0.05 Hz and 50 Hz. All recordings were sampled at 512 Hz and stored for later offline analyses. EEG recordings were then re-referenced to the contra-lateral mastoid (joint A1-A2) for the offline analyses.

### Polysomnographic (PSG) analyses

Sleep scoring and EEG pre-processing were conducted using the Deltamed coherence software (version 7.1.3.2032. Germany: Natus Europe), the PRANA software (Version 10.1. Strasbourg, France: Phitools) and Wonambi toolbox (Version 6.13; https://github.com/wonambi-python/wonambi). Two experienced raters (AAP and LB), blind to the experimental conditions, scored the different vigilance states according to the AASM recommendations^40^ (NREM 1, 2, 3, REM sleep and Wake) for each recorded night of sleep. From the scoring, we computed the duration (min) of each stage, as well as the percentage of each sleep stage relative to the total sleep period (TSP; from sleep onset to wake-up time) and relative to the total sleep time (TST; TSP minus intra-wake epochs). Latency to sleep onset (SOL, i.e., latency to first epoch of sleep) and latency to reach each stage as well as latency to reach consolidated NREM sleep (10 minutes of uninterrupted N2 and/or N3), sleep efficiency (total sleep time/time in bed*100), and number of transitions between stages were calculated. We also computed a sleep fragmentation index (SFI) as the number of shifts to wake and lighter stages (e.g., N1 or N2 from N3 or REM) divided by the total sleep time (in hours). SFI has been used previously to estimate sleep disruption^41^. Artefacts and arousals^40^ were identified visually by expert scorers (AAP and LB). Total arousal density (number/h) and per stage (number/30s-epoch for each sleep stage) were extracted.

Using the seapipe python pipeline (https://github.com/nathanecross/seapipe), the EEG spectrum power average (30 s of time resolution with artefact excluded) was calculated with a 0.2 Hz resolution, by applying a Fast Fourier Transformation (FFT; 50 % overlapping, 5 s windows, Hanning filter) using the midline frontal (Fz) and parietal (Pz) derivations. Mean power was calculated for each 0.2 bins between 0.4 and 30 Hz and for the following frequency bands: slow oscillations (0.25-1.25 Hz), delta (0.25-4 Hz), theta (4.25-7.75 Hz), alpha (8-11 Hz), sigma (11.25-16 Hz), low beta (16.25-19 Hz) and high beta (19.25-35 Hz).

#### Events detection

Spindles were detected on central channels following the procedure of Ray and colleagues (2015)^42^ and visually supervised for each night of each participant by expert scorers (AAP and LB) using the PRANA software (Version 10.1. Strasbourg, France: Phitools). We used a band-pass filtering of 8.5-12Hz for the detection of slow spindles on Fz and fast spindles were detected on Pz with a band-pass filtering of 12.5-15.5Hz^7^. Slow oscillations (SOs) were detected automatically on Fz following the procedure proposed by Staresina et al. (2015) ^43^ and implemented in the seapipe python pipeline (https://github.com/nathanecross/seapipe). SOs and spindles characteristics, including count, density (number/30s-epoch), amplitude, peak frequency and duration were extracted for N2 and N3.

#### Event co-occurrence measures and event-locked phase-amplitude coupling

Analyses of co-occurrence and phase-amplitude coupling (PAC) were conducted using the seapipe python pipeline (https://github.com/nathanecross/seapipe). A given SO and a given spindle were said to be co-occurring if they overlapped in time. To quantify the probability of the co-occurrence of SO and spindle events, we used the intersection union, a measure used in computer science and machine learning^44^. We applied a minimum threshold of 10% overlap between events, namely here whenever 10% of the duration of both events were co-occurring. Thus, within the SO-spindle complexes, we extracted the percentage of SO with spindle and SO without spindle^45^.

We also examined cross-frequency PAC between SO phases (0.5-1.25 Hz) and sigma oscillations (8.5-12Hz for Fz and 12.5-15.5Hz for Pz, as indexes of slow- and fast-spindle activity, respectively). To create the phase-amplitude distribution of slow- and fast-sigma activities, we first extracted the EEG signal from each detected event (SO) across the whole night, along with 2 seconds of buffer signal on either side of each event to avoid filter edge artefacts. We then filtered each event in the corresponding low-frequency band (0.5-1.25 Hz) and extracted the instantaneous phase time series using the angle of the Hilbert transform. In parallel, we filtered each event in their respective sigma frequency bands and extracted the instantaneous amplitude time series using the absolute value of the Hilbert transform. Next, we discarded the buffer signal around each event and binned each value in the amplitude time series by the simultaneous value in the phase (SO) time series (18 bins) to obtain the mean amplitude in each phase bin, producing a single phase-amplitude distribution per SO event. In binning mean amplitudes by phase, we divided the low-frequency cycle into 18 phase bins, balancing precision with robustness^46^.

The mean amplitudes were z-scored across phase bins for each SO event independently, to minimize the influence of amplitude differences prior to all subsequent analyses^47,48^. For each SO event, we calculated the phase bin with the maximal mean amplitude of sigma activity. The preferred coupling phase was calculated as the mean circular direction across SO events during N2 and N3. Measures of the PAC coupling strength were measured using the modulation index (MI), based on an adaptation of the Kullback-Leibler distance function for inferring statistically significant differences in the amplitude and spread of two distributions^46,49^.

#### Measures of NREM brain oscillations entrainment

The rocking bed was equipped with a sensor, which sent a marker on the EEG recording every time the bed reached the left peak of its excursion. Following a previously reported procedure^7^, we investigated whether the rocking stimulation entrained brain oscillations. In a nutshell, we computed the distribution of spindles (centre) and SOs (downstate) around the markers of rocking during the rocking night and around virtual markers (i.e., every 4 s) during the stationary night. Peri-event time histograms (PETH) provided a graphic representation of SOs and spindles around the marker (80 bins of 100 ms over the 4 s between markers).

### Questionnaires

After each experimental night, subjective sleep quality was assessed using the St Mary’s Hospital Questionnaire^50^. Specifically, we used the question “*How did you sleep last night?*” rated on a five-point scale (from very bad to very good). Within that questionnaire, participants also reported their subjective estimation of time to fall asleep (sleep latency) and total sleep duration. Sleep Perception Index (SPI) which expresses the ratio between subjective measures and objective (PSG) measures in percentage (subjective/objective*100), was calculated to assess the degree of misperception of sleep duration (SPI)^51,52^. Values around 100% indicate optimal perception while values < 100% refer to under-estimation and values > 100% refer to over-estimation of TST.

The morning after the rocking night, the pleasantness and relaxing properties of the rocking bed were evaluated by a homemade ten-point Lickert scale questionnaire (from very unpleasant/stressful to very pleasant/relaxing).

### Overnight memory assessment

To assess the impact of rocking stimulation on overnight declarative memory performance, we used a word paired-associate learning task, as we did in our previous work^7,53^, where semantically unrelated French word-pairs had to be learned. The task consisted in an encoding phase and two recall phases: an immediate (pre-sleep) and a delayed (post-sleep) recall, to assess overnight changes in memory performance. Two lists of 46 word-pairs were created, one for each experimental session. In the evening, participants were first asked to learn the 46 word-pairs presented one by one (4s each) with an inter-stimulus interval of 100 ms. Immediately after the encoding, participants underwent a first recall test during which the first word of each pair was presented, in a newly randomized order, and participants had to type the associated word (in less than 10s). In the case where participants did not remember the paired word, they were asked to guess or to leave the response blank. Feedback showing the correct associate was given at the end of each trial. After the night of sleep, a second recall test was again administered. Correct responses (hits), errors, and lack of response (misses) were measured. Memory accuracy score was computed (hits minus errors) for both recall tests and experimental sessions.

### Quantification & Statistical Analysis

Statistical analyses were conducted using RStudio 1.2.50 (RStudio, Inc., Boston, MA) and R custom scripts and functions (e.g., packages nparLD, rstatix, car).

Normality of data distribution was checked with Shapiro tests and homogeneity of variance was tested with Levene tests. When normal and homogeneous, we used mainly paired t-tests and repeated-measures analyses of variances (ANOVA), with post hoc multiple comparison tests (using Bonferroni correction) to specify main and/or interaction effects whenever needed. When variance was not normal and/or homogeneous, we used non-parametric tests (paired Wilcox Test or Wald-Type Statistic). All tests included a repeated measure factor Condition (rocking, stationary). We computed effect sizes to indicate the degree of change in response to Condition using Hedges’s *g* (corrects for small sample size). Concerning rocking entrainment, chi-square goodness of fit test was used to assess the non-uniformity of the occurrence of SOs and spindles in N2 and N3 averaged across participants in each condition. A 4s cycle (period of 4s post rocking marker) was used to measure chi-square goodness of fit against the null hypothesis that the distribution of events is uniform across bins (i.e., the probability of the occurrence is equal in all bins) around the markers. Degrees of freedom were corrected according to Greenhouse-Geisser when necessary. The level of significance was set to p-values <.05. All statistical analyses concerning sleep architecture were done on the 16 subjects included in this study. Due to technical issues, the rocking markers were not available for 3 participants. Therefore, this analysis was performed on 13 participants. Due to a MATLAB issue during the cognitive test, the data from one participant was lost and the statistical analysis of memory performance was performed on the data from 15 participants. Exploratory Pearsons’ correlations were conducted between changes in sleep variables.

## RESULTS

Sixteen individuals presenting with insomnia symptoms and objective markers of poor sleep participated in this crossover design. Participants were young adults (24 ± 3 years old; 11 female) suffering from chronic insomnia with an ISI score >8 (mean score 14.8 ± 2.7; **Table 1**).

### Self-reported sleep quality and perception of rocking

Based on the participants’ responses to the St Mary’s Questionnaire in the morning after each experimental night, we found that participants reported having a better sleep after the rocking night compared to the stationary night (+25%, *p*=.008, *g’*=-0.72; **Figure 2A**), while the degree of sleep misperception did not differ between the conditions (SPI-rocking: 93.8±11.5%; SPI-stationary: 94.7±13.6%; *p*>.69, *g’*<0.1; **Figure S1**).

**Figure 2.**
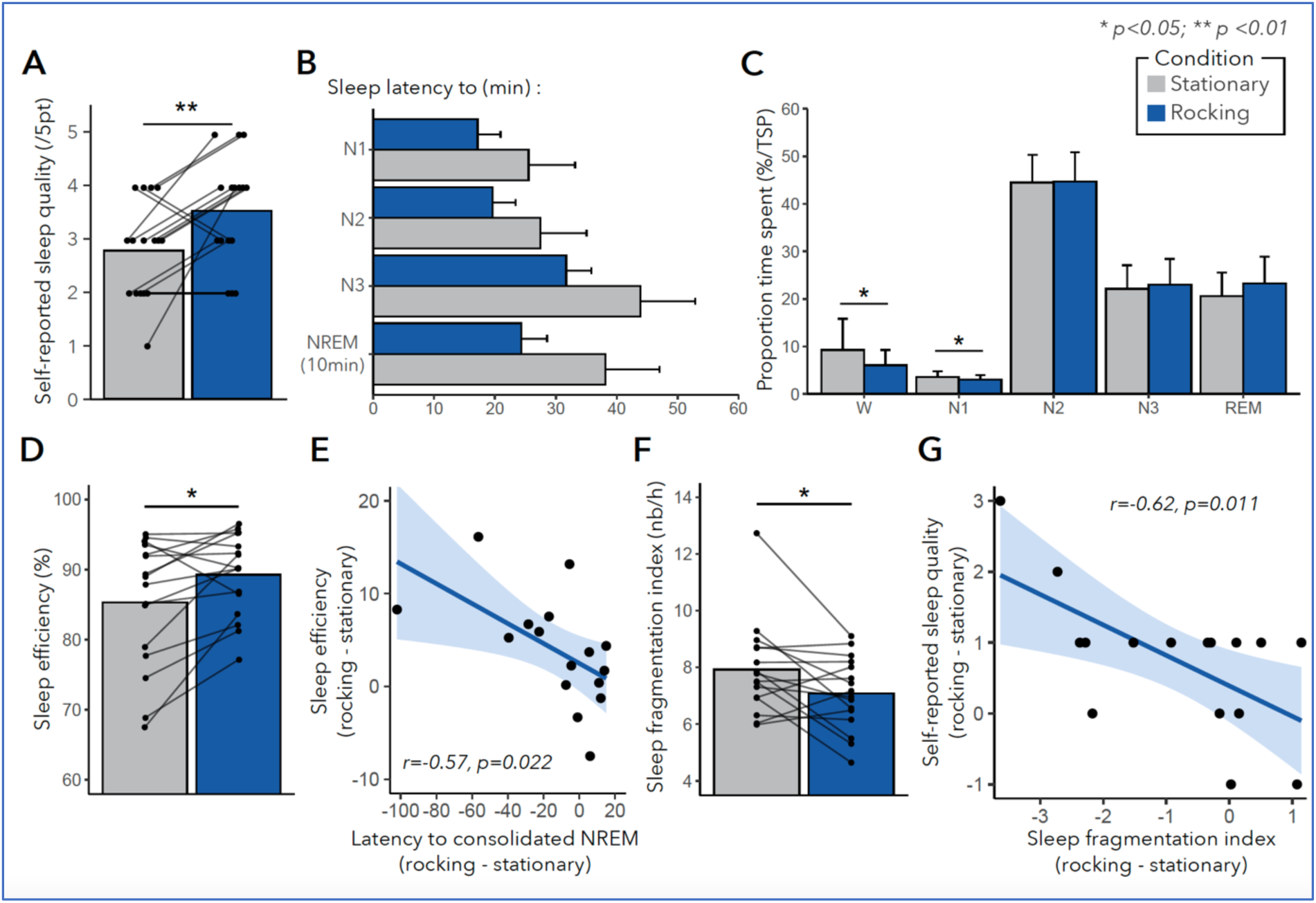
Effects of rocking on sleep architecture. (A) Mean and individual-specific self-reported sleep quality after stationary (grey) and rocking (blue) nights. Scale of 5-point from 0 = very bad sleep to 5 = very good sleep (B) Mean (±SD) sleep latencies (in min) to N1, N2, N3 and to consolidated NREM (i.e., at least 10 min of uninterrupted NREM sleep) during stationary and rocking nights (C) Mean (±SD) sleep stage distribution (percentage of total sleep period) during stationary and rocking nights (D Mean and individual-specific sleep efficiency (in percentage) during stationary and rocking nights (E) Scatter plot showing a significant correlation between the change (rocking minus stationary) in sleep latency to consolidated NREM (at least 10min of uninterrupted NREM) and sleep efficiency (F) Mean and individual-specific sleep fragmentation index (number per hour) during stationary and rocking nights (G) Scatter plot showing a significant correlation between the change (rocking minus stationary) in sleep fragmentation index and self-reported sleep quality * p<.05 ** p<.01, N=16

Using a ten-point Likert scale questionnaire on rocking pleasantness and relaxing properties, poor sleepers reported being generally satisfied by the rocking stimulation (7.8 ± 1.2/10). While the level of pleasantness or relaxation due to the rocking did not correlate with any objective change in sleep (e.g., wake duration, spindle density; all *p*>.05), we found that individuals who exhibited higher anxiety (anxiety subscale of the HADS questionnaire) and higher sleep complaints (PSQI) reported the rocking stimulation as more pleasant/relaxing (*r*=0.56, *p*=.03 *and r*=0.58, *p*=.02, respectively).

### Rocking stimulation decreases time spent awake and sleep fragmentation in poor sleepers

Compared to the stationary night, we found no significant effect of rocking on sleep onset latency (SOL; *p*=.67; *g’*=0.31) or latency to 10 minutes of consolidated NREM sleep (*p*=.19; *g’*=0.42). Moreover, rocking did not affect latencies to the different sleep stages (all *p*>.05; *g’* from 0.2 for SL to N2 and REM to 0.37 for SL to N3; **Figure 2B**).

While there was no effect of rocking on total sleep time (*p>*.05), we assessed the effect of rocking on time spent in each stage (percentage of TSP). We found an interaction Condition by Stage (*S*=11.9, *d.f*.=4, *p*=.017) driven by a significant decrease in time spent awake (-34%; *p*=.015, *g’*=0.53) and time spent in N1 (-15.5%; *p*=.038, *g’*=0.48). Time spent in N2, N3 and REM remained largely unchanged (all *p*>.05; all *g’*<0.2; **Figure 2C**). Yet, the decrease in time spent awake (%TSP) was associated with extended time spent in the deeper stages of NREM (N2+N3; *r*=-0.55, *p*=.027) but not REM duration (*r*=-0.47, *p*=.063). As a result of these stage duration changes, the reduction in wake duration was associated with increased sleep efficiency (SE; *r*=-0.73; *p*=.0013), with a 4.5% (*p*=0.016*, g’*=-0.42) increase in sleep efficiency during rocking compared to stationary night (**Figure 2D**). The increase in SE was also associated with a faster entrance in consolidated NREM sleep (*r*=-0.57, *p*=.022; **Figure 2E**).

While no significant changes in N2 and N3 were observed, individuals seemed to differ in the modulation of N2 and/or N3 sleep stages in response to rocking stimulation, whereby 5/16 participants (31.2%) had an increase in N3 only, 6/16 (37.5%) in N2 only, and 5/16 (31.2%) with an increase in both N2+N3.

Concerning measures of sleep fragmentation, we found no significant effect of rocking on the density of arousals overnight (*p*=.66, *g’*<0.1) or per stage (all *p*>.05). However, we found a reduction in the sleep fragmentation index (SFI) during the rocking night compared to stationary (*p*=.034, *g’*=0.53; **Figure 2F**). Interestingly, such a reduction in the sleep fragmentation index correlated with higher subjective sleep quality (*r*=-0.62, *p*=.011; **Figure 2G**).

### Rocking stimulation impacts NREM brain oscillations

When investigating slow (Fz; 8.5-12.5Hz) and fast (Pz; 12.5-15.5Hz) spindles, poor sleepers exhibited a boost in fast spindles density during N3 when rocked (*p*=.003, *g’*=-0.33; **Figure 3AB**). However, there was no change in slow spindles density (*p*=.97, *g’*<0.1) and no change in N2 for both slow and fast spindles, as well as no change in spindle characteristics (i.e., amplitude, frequency, duration; **Figure 3A**) or sigma spectral activity (11.25-16Hz) during N2 and N3 (all comparisons *p*>.05). We also found no change in slow oscillation (<1Hz on Fz) activity (i.e., density, count, amplitude, peak frequency) in both N2 and N3 (all *p*>0.1; **Figure 4A**).

**Figure 3.**
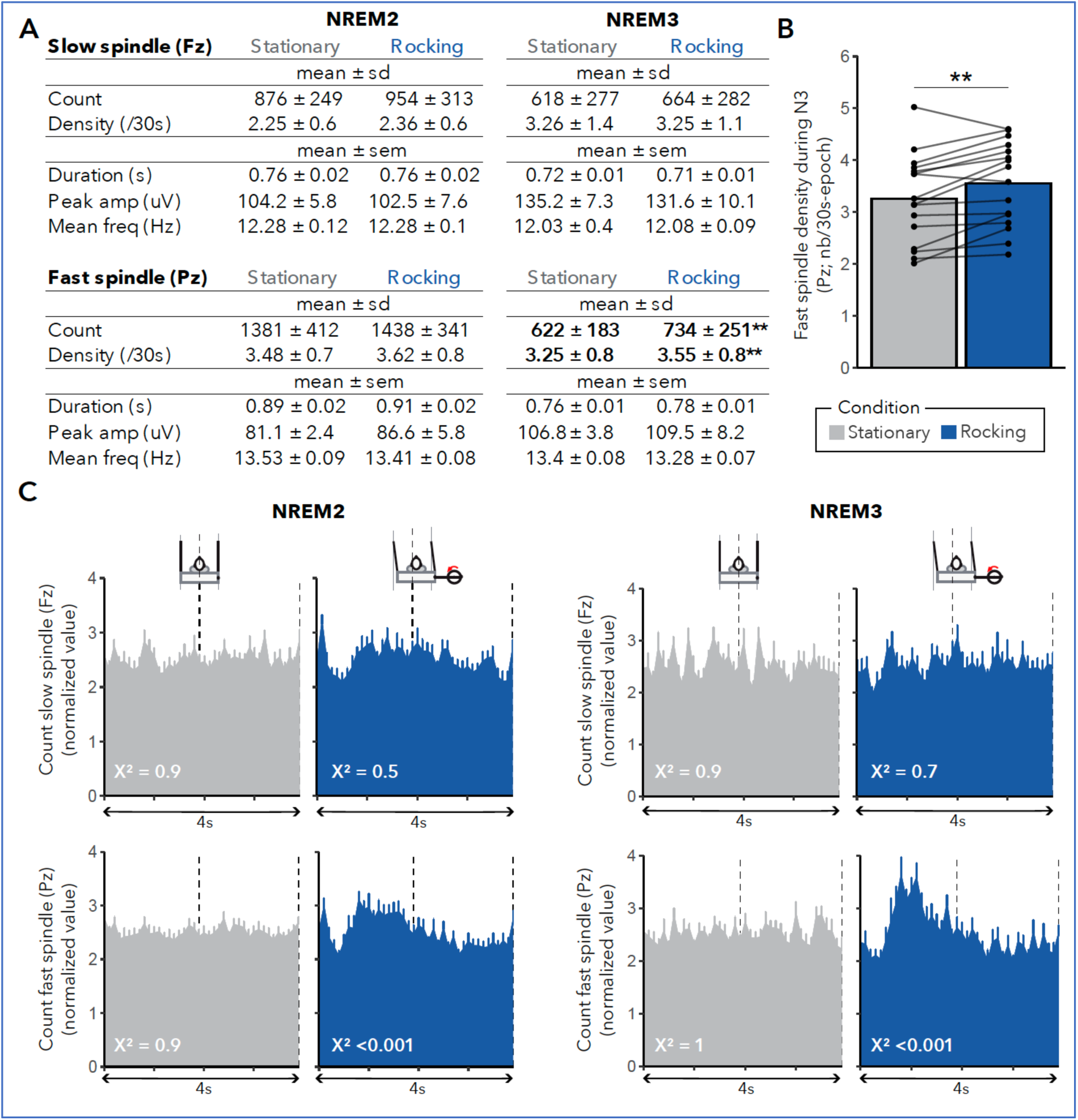
Effects of rocking on spindle activity. (A) Mean (±SD) slow and fast spindles count and density and mean (±SEM) for spindles characteristics (duration, peak amplitude, mean frequency) recorded from Fz (top) and Pz (bottom) in N2 (left) and N3 (right) during stationary and rocking nights (N=16) (B) Mean and individual-specific density of fast spindles (12.5-15.5Hz, detected on Pz) in N3 during stationary (grey) and rocking (blue) nights (N=16) (C) Peri-event time histograms (PETHs) of mean (±SEM) count of slow spindles (detected on Fz - top) and fast spindles (detected on Pz - bottom) after the rocking marker (at each left turning point of the movement; time scale of 4 s, 80 bins of 100 ms) in N2 (left) and N3 (right) during stationary and rocking nights across 13 participants.

**Figure 4.**
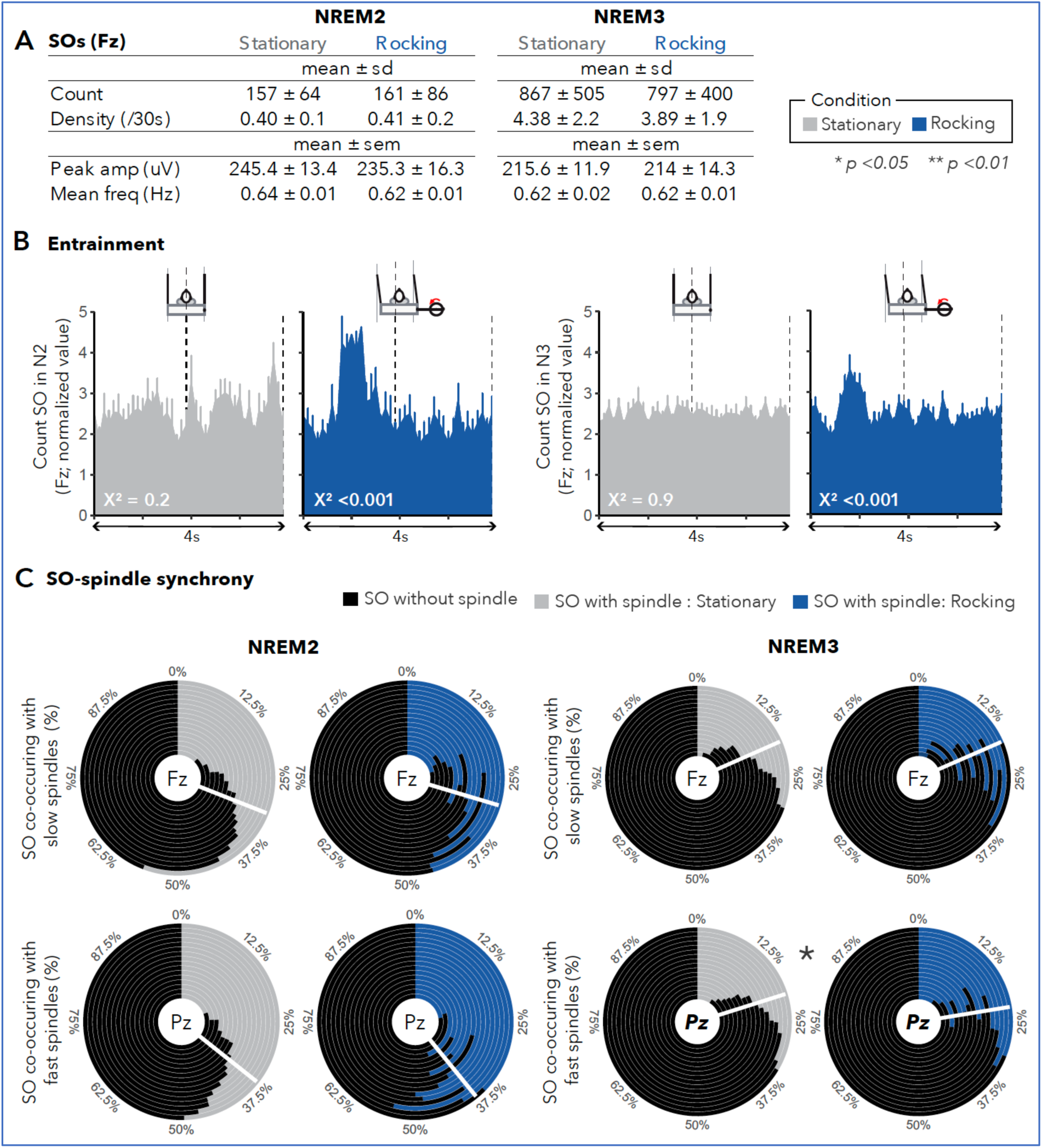
Effects of rocking on slow oscillations (SOs) (A) Mean (±SD) count and density of slow oscillations (SO) and mean (±SEM) characteristics for SOs (peak amplitude, frequency) recorded from Fz during N2 (left) and N3 (right) during stationary and rocking nights (N=16) (B) Peri-event time histograms (PETHs) of mean (±SEM) count of SOs (detected on Fz) after the rocking marker (i.e., at each left turning point of the movement; time scale of 4 s, 80 bins of 100 ms) in N2 (left) and N3 (right) during stationary (grey) and rocking (blue) nights across 13 participants. (C) Racetrack plot of percentage of SO-slow spindles (detected on Fz – top) and SO-fast spindles (detected on Pz – bottom) co-occurrence (in grey/blue) and percentage of SOs without spindles co-occurring (in black) in N2 (left) and N3 (right) during stationary and rocking nights per individuals (N=16). The white bars represent the mean percentage of SO-spindle co-occurrence per night.

Regardless of the impact of rocking on SOs and sleep spindles density, we observed that the occurrence of both SOs and spindles clustered at specific time points with respect to the rhythmic motion of the bed (see Methods for details; **Figure 3C and Figure 4B**), suggesting entrainment of both spindles and SOs during NREM sleep at each turning point of the bed (i.e., maximal linear acceleration). Statistical analyses confirmed that, in the rocking condition, the distributions of fast spindle (**Figure 3C**) and SOs (**Figure 4B**) were not uniform during N2 and N3 (chi-square *p*<.05), while the distribution of slow spindles was uniform (*p*>.05; **Figure 3C**). During the stationary night, the distribution of SOs, slow and fast spindles around imposed markers (i.e., every 4s) were uniform during N2 and N3 (all chi-square *p*>0.1; **Figure 3C and Figure 4B**).

Next, we analysed and compared spindle-SO coupling characteristics between rocking and stationary conditions and found that the proportion of SO occurring with a fast spindle (Pz) during N3 increased when rocked (+11%; *p*=.044; **Figure 4C**). Conversely, no change was observed for the SO occurring with fast spindles (Pz) in N2 and for slow spindle (Fz) in N2 and N3 (all *p*>.05; **Figure 4C**). We found no change in the preferred phase or magnitude of SO-sigma modulation (all *p*>.05; **Figure S2**), suggesting that the mechanisms underlying the specific temporal relationship between co-occurring spindles and SOs were not affected by rocking.

### No effect of rocking stimulation on memory consolidation in poor sleepers

We found no difference in overnight change (morning minus evening) in memory accuracy (correct minus errors) between stationary and rocking nights (N=15; *p*>0.1, *g’*<0.1; **Figure S3A**). Moreover, repeated measure ANOVAs revealed no effect of Condition (*F*(1,14)=0.046, *p*=.83), Session (*F*(1,14)=0.453, *p*=.052) or interaction Condition by Session (*F*(1,14)=0.005, *p*=.94) on memory accuracy. Similar results were found for correct responses, errors and misses (all *p*>0.05, **Figure S3B**).

## DISCUSSION

We previously showed that gentle rocking stimulation (0.25Hz - lateral) during a whole night of sleep can improve sleep quality (i.e., shorter sleep onset, deeper sleep, less fragmentation) in good sleepers^6,7^. It has been shown that rocking stimulation requires optimal settings to be beneficial for sleep^8,54–56^. Specifically, it appears that linear lateral motion may be the perfect candidate to improve sleep^6,7,57,58^. By specifically targeting sleep difficulties reported by clinical populations, sleeping in a rocking bed could become a relevant sleep intervention in individuals with sleep difficulties such as insomnia. Insomnia is a sleep disorder whose clinical diagnosis criteria are solely based on subjective sleep complaints^10,11^ not always corroborated by objective measures, such as polysomnographic (PSG) recordings^59^, which makes this complex disorder challenging to investigate, define and manage^25,27,28,60–62^. Here, to avoid some of this complexity, we deliberately selected a sample of young individuals reporting insomnia symptoms who also presented markers of objective poor sleep during 3-weeks of actigraphy recording and one screening PSG recording in-lab. As such, our sample represents a subtype of chronic insomnia characterized by subjective complaints (ISI >8)^34^ and an average sleep efficiency below 85%^19,39^ (i.e., previously categorized as suffering from psychophysiological insomnia; ICSD-2^30,31^).

Compared to a typical stationary night, we found that rocking reduced the time spent awake and in light sleep (N1). In line with those changes, we found an increase in objective sleep efficiency and a decrease in sleep fragmentation index^41^, suggesting less switching from deeper sleep to light sleep or wake. Critically, given the participants’ primary complaints, these changes in sleep were paralleled by a significant improvement in the subjective assessment of the quality of their sleep. Further corroborating this link, reduced sleep fragmentation index correlated with higher subjective sleep quality. Finally, gentle rocking stimulation boosted fast spindle activity during N3 and entrained NREM brain oscillations with a rhythmic appearance synchronized to the bed movement, which is consistent with our previous work on good sleepers^6,7^.

Insomnia diagnosis criteria being solely based on subjective reports^10,11^, it is clinically relevant to see a better satisfaction of their sleep quality after rocking compared to a stationary night. While we only tested the effects of a single night of rocking, we found a large effect size on self-reported sleep quality (*g’*= 0.72) resembling those reported in studies investigating the benefit of CBTi, the first line of treatment for chronic insomnia^22,23^. These results suggest a promising new avenue of research for alternative treatments for the management of insomnia. Yet, whether sleep satisfaction can remain high and stable over multiple nights of rocking, thus recommended for daily use at-home, remains to be tested.

In our sample, rocking led to objective changes in sleep that directly address one of the main complaints of insomnia, i.e., difficulty maintaining sleep throughout the night^10,11^. Indeed, during the rocking night, individuals exhibited a reduction in time spent awake and in lighter sleep, yielding an increase in sleep efficiency as well as a reduction in sleep fragmentation index (SFI) and suggesting enhanced sleep maintenance^41^. Interestingly, the change in SFI was associated with an increase in sleep satisfaction after a rocking night. This is consistent with previous work showing that, compared to traditional macro-architecture parameters, poor sleep stability (state-transition frequency) is a key determinant of subjective sleep quality^63^. Of note, we did not observe any change in the degree of sleep misperception, suggesting that the change in the subjective rating of sleep is not due to a change in sleep perception^21^ but rather to the objective change in sleep.

Overall, these results are in line with our previous work on good sleepers^7^ and rodents^8^ where we argued that the rocking mechanism originates from the vestibular receptors (otoliths), which send the information of continuous and rhythmic movement to the thalamus, thus providing a synchronizing influence on neuronal activity within thalamocortical networks. The use of a rocking bed might thus be an intervention of choice to aid individuals complaining of long awakenings in the middle of the night^10,11^.

Complaints about falling asleep are also extremely common in insomnia and it has been previously demonstrated that rocking accelerated entrance into deeper sleep in good sleepers^6,7,55,58^. However, in poor sleepers, while in the rocking condition, 43.7% (7/16) of participants had shorter SOL and 62.5% (10/16) participants reached consolidated NREM sleep faster, these changes in SOL or latencies to different stages did not reach statistical significance. While the rocking-related reduction in time spent awake and in lighter stage appeared common to our sample, latencies to sleep and time spent on NREM stages showed large interindividual variations (which we did not statistically assess due to power limitation), unlike good sleepers who exhibited a homogeneous boost in N3 during the rocking night^7^. The heterogeneity in response to rocking stimulation in individuals with insomnia may reflect the complexity of the disorder, which encompasses multiple profiles of symptoms and etiologies^25,27,28^. Specifically, our sample showed mixed effects of rocking on N2 and N3 duration, with some participants having similar patterns to good sleepers (i.e., more N3, less N2) while others exhibited increases in N2 or both N2 and N3. Another possible mechanism underlying the heterogeneity in response to rocking in poor sleepers may not relate to sleep processes per se but to variations in sensory processing^64^. Individuals with insomnia are more prone to high sensory reactivity^65^, characterised by greater depth of information processing and awareness of environmental stimuli^66^. For instance, recent studies using external stimulation involving tactile and proprioceptive inputs (e.g., weighted blanket) have been shown to reduce insomnia severity^67,68^. If individuals with poor sleep exhibit variable levels of sensory-processing sensitivity, it may explain why they responded differently to the sensory stimulation induced by the rocking bed. Larger future clinical trials on rocking stimulation on multiple stimulation nights might reveal whether insomnia subtype-specific disturbances, based on specific insomnia symptom patterns, objective sleep alterations^28^ (as done here), presence of sleep misperception^51,69^, or psychological traits^25^, respond more strongly to rocking compared to others.

We previously reported that rocking enhanced overnight consolidation of declarative memory in good sleepers, and suggested that this effect was plausibly mediated by the increase in spindle activity (in particular during N3) and enhanced spindles and SOs synchronization with the rocking periodicity^7,9^, in line with the proposed role of temporal phase binding of cortical (SOs), thalamic (fast spindles), and hippocampal (sharp wave ripples) rhythms during NREM sleep in declarative memory consolidation processes^70,71^. In the present study in poor sleepers, we also observed no evidence for enhanced memory consolidation during rocking despite a boost in fast spindles and similar entrainment processes, particularly for SOs and fast spindles in N2 and N3. While these fundamental thalamocortical oscillatory mechanisms seemed to be similarly influenced in good and poor sleepers, we may speculate that the absence of memory effects in poor sleepers might rather relate to the variable impact of rocking on the duration of N2 and N3 sleep stages. Yet, and as mentioned above, accounting for the effects of rocking on memory in subgroups of participants would require larger sample sizes.

In summary, the present findings demonstrate that applying rhythmic sensory stimulation during a whole night of sleep boosts sleep maintenance (i.e., decreases in sleep fragmentation and light sleep duration), with an associated increase in sleep efficiency, and improves subjective sleep quality in individuals with insomnia complaints and objective poor sleep. In other words, sleeping on a rocking bed could be considered an alternative or complementary intervention for the management of insomnia in poor sleepers. Whether the beneficial effect of rocking bed could persist over multiple nights and reduce insomnia severity, including daytime functioning complaints^32^ in the long term, remains to be tested in a randomized controlled study in a large sample of individuals with chronic insomnia.

## ACKNOWLEDGEMENT

This work was supported by grants from the Swiss National Science Foundation (CR3113_149731; 320030_182589; 320030_159862). We thank the staff from the Center for Sleep Medicine (HUG) as well as Ilde Pieroni for their help in data collection.

The authors declare no competing interests.

## AUTHOR CONTRIBUTIONS STATEMENT

*Aurore A. Perrault*: Conceptualization, Project administration, Investigation, Data curation, Formal analysis, Methodology, Visualization, Interpretation, Writing – original draft, review & editing

*Nathan E. Cross:* Methodology, Interpretation, Writing – review & editing

*Thien Thanh Dang Vu*: Writing – review & editing

*Sophie Schwartz*: Conceptualization, Funding acquisition, Supervision, Interpretation, Writing – original draft, review & editing

*Laurence Bayer:* Conceptualization, Funding acquisition, Project administration, Investigation Data curation, Supervision, Interpretation, Writing – original draft, Writing – review & editing

