## Supplemental Material for "Whole-night gentle rocking improves sleep in poor sleepers with insomnia complaints"

### Supplemental results

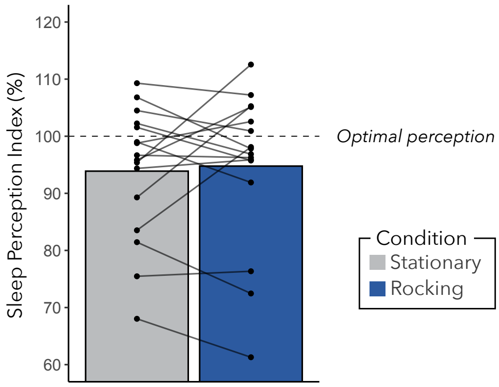


**Figure S1 – Degree of sleep misperception**

*Mean and individual-specific sleep perception index (in%) during stationary (grey) and rocking (blue) nights (N=16)*

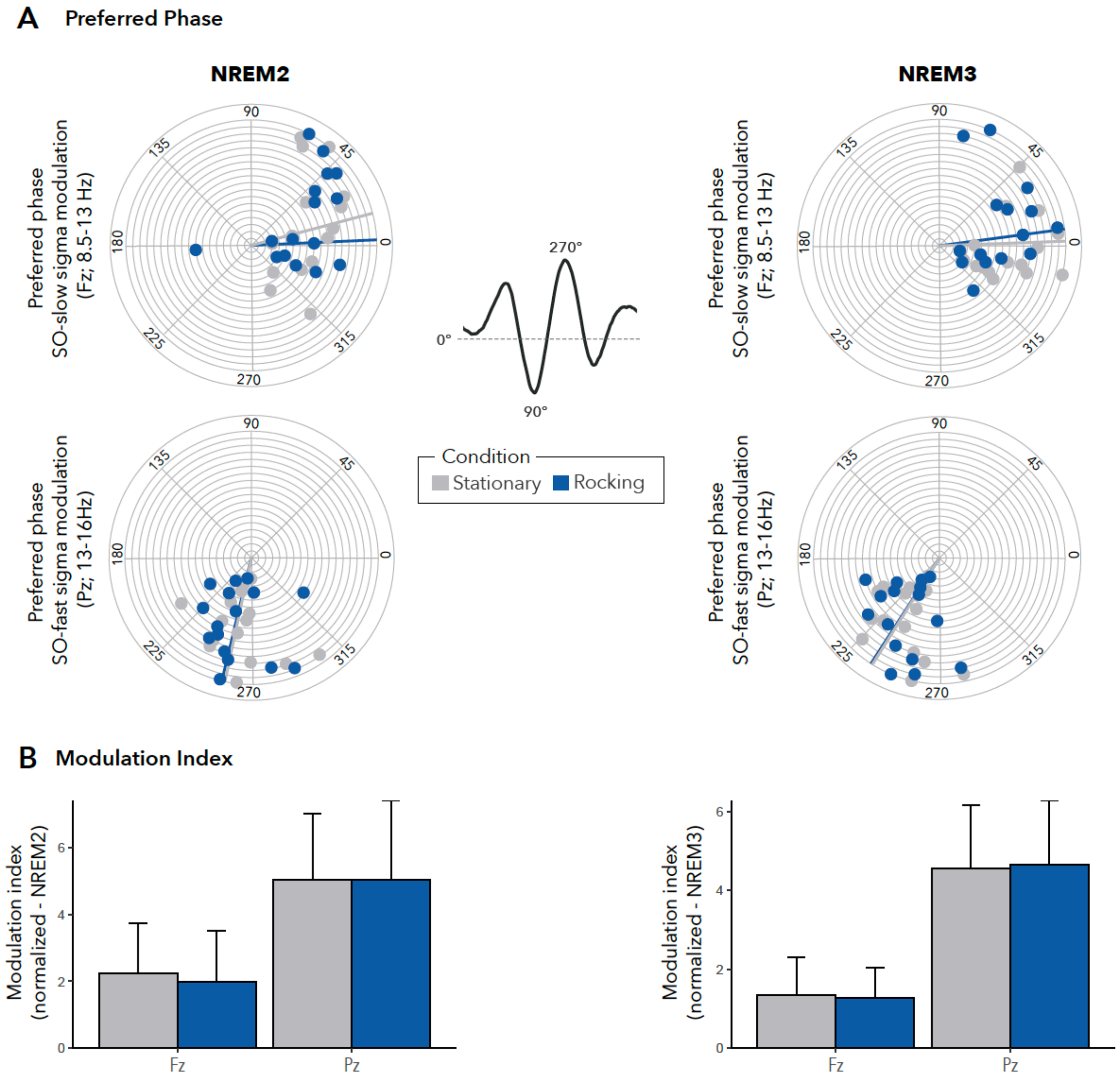


**Figure S2 – No effects of rocking on phase-frequency coupling between SO and spindle activity**

*(A) Unit circle of preferred phase of the SO-slow sigma power modulation (Fz; top) and SO-fast sigma power (Pz; bottom) for NREM2 (left) and NREM3 (right) for each participant during stationary (grey) and rocking (blue) nights.*

*Each line represents a participant and contains 2 small circles reflecting preferred phase during stationary and rocking nights.*

*(B) Mean (±SD) modulation index of SO-slow sigma power modulation (Fz) and SO-fast sigma power (Pz) for NREM2 (left) and NREM3 (right) during stationary and rocking nights*

A

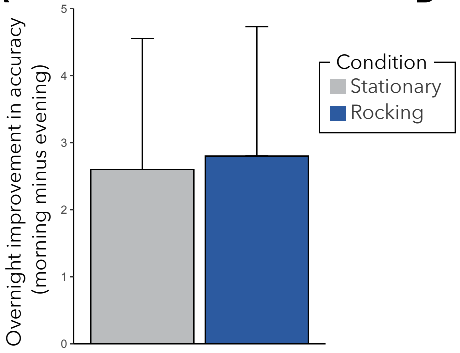


B

Stationary (mean ± sd)

| | Evening | Morning | *p value* |
|---|---|---|---|
| Number of hits | 14.6 ± 11.22 | 16.4 ± 13 | *0.15* |
| Number of errors | 9 ± 11.98 | 8.13 ± 11.24 | *0.39* |
| Numbers of misses | 22.33 ± 12.45 | 21.4 ± 13.2 | *0.58* |

Rocking (mean ± sd)

| | Evening | Morning | *p value* |
|---|---|---|---|
| Number of hits | 16 ± 11.5 | 17.6 ± 12 | *0.19* |
| Number of errors | 9.86 ± 9.78 | 8.66 ± 10.64 | *0.38* |
| Numbers of misses | 20.13 ± 11.45 | 19.7 ± 13.6 | *0.74* |

Figure S3 – No effects of rocking on overnight memory performance

*(A) Mean (±SD) overnight improvement in accuracy (hits minus errors) during stationary (grey) and rocking (blue) nights.*

*(B) Mean (±SD) number of hits, errors and misses performed during the evening and morning recalls for stationary and rocking nights.*
